# Minimization of muscle activation costs demanded from mechanical work and power accounts for selection of duty factor in human gaits

**DOI:** 10.1101/205526

**Authors:** Grzegorz Sobota, James R. Usherwood

## Abstract

Duty factor DF – the proportion of a stride a foot is in contact with the ground – is of fundamental mechanical importance, and is often viewed as a defining kinematic parameter distinguishing walking (DF>0.5) from running (DF<0.5). However, the mechanical and/or physiological considerations that determine duty factor are not well understood. Here, a model is proposed that focuses on the interaction between mechanical and muscle costs to account for duty factor in human gaits. It minimizes the activation costs associated with mechanical work or power demand during muscle contraction (whichever is the more demanding). Empirical observations match model predictions using initial muscle parameters over a range of speeds within gaits. However, a better match is achieved – and a better account for the walk-run transition – with tuned muscle parameters. The tuned model is validated with responses in duty factor to walking at a range of imposed, unnatural step frequencies.

## Notations

| Term | Definition or description |
| --- | --- |
| BM | Body mass |
| BW | Body weight |
| $C_{MA}$ | Cost of muscle activation |
| CT | Cost of Transport |
| $CT_P$ | Cost of Transport due to muscle activation demanded for $mP_{cc}$ |
| $CT_W$ | Cost of Transport due to muscle activation demanded for $W_{cc}$ |
| $CT^F$ | Cost of Transport with constant frequency assumption |
| $CT^{Tsw}$ | Cost of Transport with constant swing time assumption |
| DF | Duty Factor: the proportion of the stride cycle a foot (left or right) is in contact with the ground |
| F | Step frequency |
| g | The magnitude of gravitational acceleration ($9.81 \text{ ms}^{-2}$) |
| GRF | Ground Reaction Force |
| $kg_{am}$ | Kilograms of activated muscle mass |
| $kg_{bm}$ | Kilograms of body mass |
| L | Leg length |
| $mP_{cc}$ | Mean mechanical power over the duration of concentric contraction |
| $P^*$ | Muscle mass specific power capacity |
| SL | Step length |
| $T_{sw}$ | Swing (protraction) duration |
| $V_{max}$ | Maximum muscle velocity (in lengths / s) |
| $V_n$ | Non-dimensional velocity (the square root of the prevalent form of Froude number in biomechanics). |
| $W_{cc}$ | Mechanical work of concentric contraction (positive) |
| $W^*$ | Muscle mass specific work capacity |
| Additional suffixes |  |
| Walk | Relating to walking |
| Run | Relating to running |

## Introduction

A great deal of research has focused on energy expenditure in the area of human locomotion. Much of this analysis has been related to the factors influencing the metabolic cost of transport (metabolic cost per distance travelled, Zarrugh et al., 1974). It is well accepted that, under free locomotion conditions, selected human gait patterns are consistent with minimum metabolic cost (Sparrow and Newell, 1998; Alexander, 2002). To walk or run at a certain speed, humans have the choice of an infinite number of combinations of frequency and step length (Bertram and Ruina, 2001), which may determine other spatio-temporal parameters. Previous studies do not indicate clearly the dependence of running cost of transport on speed (Margaria et al., 1963; Mayhew, 1977; Steudel-Numbers and Wall-Scheffler, 2009), but for a given speed, each subject selects only one specific set of kinematic variables resulting in low transport costs compared to other possible values (Högberg, 1952; Gutmann et al. 2006). Deviations from the preferred step length, width and frequency result in increased energy expenditure for walking (Zarrugh and Radcliffe, 1978; Donelan et al., 2001; Kuo et al., 2005; Selinger et al., 2015). An additional factor is practice, which decreases level of muscle activation and metabolic energy expenditure (Komi et al., 1987; Strasser and Ernst, 1992; Lay et al., 2002). However, the mechanical and physiological factors that determine the metabolic costs of walking and running remain unclear.

A metabolic cost to muscle activation - separate from that associated directly with cross-bridge cycling - can be demonstrated with *in vitro* heat measurements of isometric muscles held at a range of lengths (Homsher et al., 1972). Heat is released even when a muscle is held at a sufficient length that cross-bridges barely overlap (though the muscle is undamaged), and there is minimal tension. This energetic cost ‘reflects the energetics of calcium release and reaccumulation’ (Homsher et al., 1972). Barclay (2012) supports this mechanistic account for a metabolic cost of activation associated with pumping ions for a contraction. Might this metabolic demand provide insight into the costs determining economical gait parameters?

For a range of gait features there is empirical evidence that metabolic cost minimisation relates closely to activated muscle volume during level legged locomotion (Bertram and Ruina, 2001; Donelan et al., 2001). Furthermore, ‘Cost of muscle force’ models applied to a range of animals are effective at relating metabolic costs to the costs of activating muscle to meet imposed forces (Taylor, 1985, Kram and Taylor, 1990; Roberts et al., 1998; Doke and Kuo, 2007; Pontzer et al., 2009; Pontzer, 2016). Hubel and Usherwood (2015) propose that the fundamental requirements for muscle loading, activation and therefore cost of locomotion are mechanical work and power during contraction, given a limited capacity for a volume or mass of muscle to produce positive work and power. The difference from previous approaches is that a ‘cost of muscle force’ is not included in its own right. They assumed that only those forces that are required for the work and power demands are applied to the muscle. This is based on the assumption that the Effective Mechanical Advantage or Gear Ratio that relates Ground Reaction Forces to muscle forces can be appropriately tuned, in evolutionary terms through bone and tendon geometry, or behaviourally through small adjustments to posture with GRFs passing close to joint centers. Using extremely reductionist models, many aspects of gait kinematics and kinetics have been found to scale with speed and size in a manner that is consistent with minimising the muscle activation required for the more demanding between mechanical work and power (Hubel and Usherwood, 2015). This approach assumes that the metabolic costs associated with activation dominate; that cross-bridge cycling costs to perform work are negligible. This is clearly not applicable in cases where there are large net work demands - such as in incline locomotion (e.g. Pontzer, 2016).

Here, we expand the reductionist models to explore whether minimization of costs of activation alone can account for, and predict, further aspects of human gait. Specifically, duty factor (DF) - the ratio of foot contact duration with the ground to the total cycle - is of interest, not least because it presents the kinematic boundary between walking (DF>0.5) and running (DF<0.5). Computer optimization to find mechanical work-minimizing gaits - at least for a point-mass reduction - finds walking at low speeds at a duty factor of exactly 0.5, and near-impulsive running, with duty factors approaching zero (Srinivasan and Ruina, 2006). However, these ‘work-minimising’ gaits also demand momentary periods of infinite mechanical power. Can simple considerations of the costs of muscle action - which are influenced by both work and power demands - account for the duty factors actually observed in human walking and running? To approach this question, we use models that allow us to combine the influences of speed and DF on ground reaction forces in order to predict the energetic implications of a kinematic parameter space. Empirical measurements are made in order to provide a range of necessary input parameters, and to validate simulation results.

## Methods

### *Methods* overview

Our approach is to calculate the mechanical implications of duty factor with simple models for walking and running ground reaction forces (fig. 1). Assumptions are required concerning protraction (swing) timing; a couple of options are considered. From the model forces and dynamics, mechanical work and power requirements are calculated, and their cost implications are determined in terms of muscle activation demands. Activation cost surfaces (depending on speed and duty factor) are initially calculated based on muscle properties assumed in previous studies (Hubel and Usherwood, 2015), justified only as being ‘empirically reasonable’. Subsequent analyses allow tuning of muscle properties to best match kinematic observations; these are then validated against measurements of walking at a range of unnatural, imposed step frequencies (step lengths).

**Figure 1.**
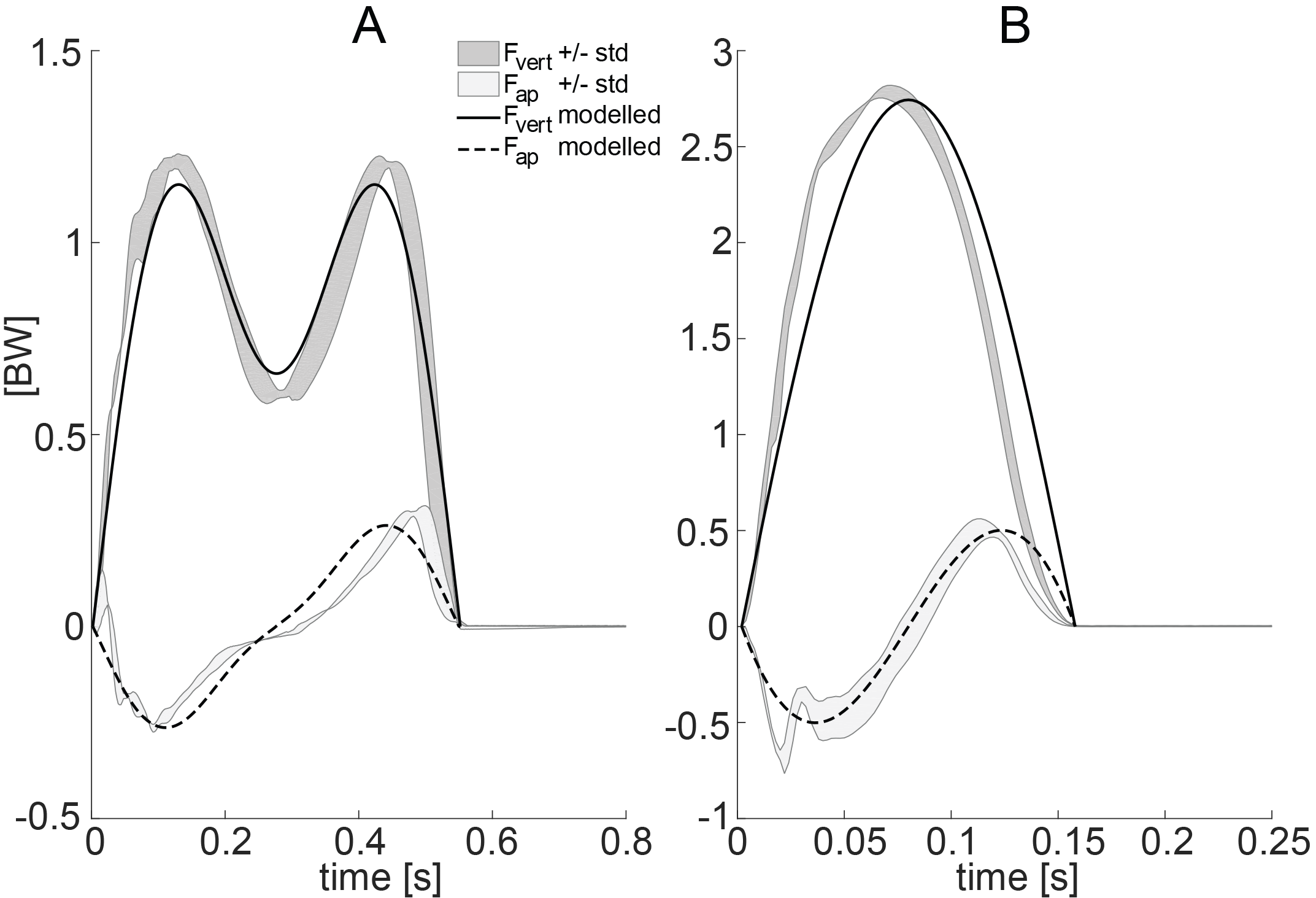
Examples of ground reaction forces (BW – body weight) for walking (A, 0.6 V_n_) and running (B, 2.1 V_n_) of experimental data (grey regions showing +/− 1SD [BW]) and simulation (black lines). Dark grey and solid lines denote vertical forces and light grey and dot-dashed lines anterio-posterior forces. Such and similar models provide a good fit for measured force data for adult humans over a range of speeds in humans (Hubel and Usherwood, 2015) and running (Robilliard and Wilson, 2005; Srinivasan and Holmes, 2008).

### Vertical ground reaction force model: walking

Model ground reaction forces acting along the limb for walking were determined from the addition of two sine waves describing vertical ground reaction forces, largely following Alexander and Jayes (1980) (fig. 1A). The amplitudes of the two sine waves can be derived analytically for a given duty factor, leg length and velocity (Hubel and Usherwood, 2015) with the assumption that midstance forces match those of a stiff-limbed vaulter (justified as being consistent with a pure work-minimizing gait, and matching observation in adult humans over a range of speeds (Usherwood et al., 2012)). Walking speeds are not achievable above a normalised velocity (V_n_=V·(L·g)^−1/2^ where V is forward speed, L is limb length and g=9.81 ms^−2^) of 1: above this, midstance forces for a stiff-limbed vault fall below zero, and leg-tension would be required (Alexander, 1980).

### Vertical ground reaction force model: running

For running, the following kinematic data were used: time of flight and necessary take off vertical velocity. The shape of the vertical ground reaction force curve was a half-sine wave (Alexander and Jayes, 1980) that meets the impulse requirements during stance to, averaged over a step, support body weight (fig. 1B).

### Horizontal ground reaction forces

Antero-posterior (fore-aft) forces were calculated from the vertical forces and the assumption that forces act in line with the leg, and the top of the leg approximates the centre of mass. While this is wrong in detail (and measurable torques exist about the centre of mass during stance in running may have some work-reducing role – Usherwood and Hubel, 2012), this provides a good initial approximation to the dominating energy fluctuations. In running, the analytical sine-wave approach used here approximates numerical spring-mass (sometimes termed Spring Loaded Inverted Pendulum or SLIP) models closely (Robilliard and Wilson, 2005; Daley and Usherwood, 2010). Figure 2 shows model-derived work and power demands for self-selected walking and running at a range of speeds, and compares these with empirical measurements derived through inverse dynamics.

**Figure 2.**
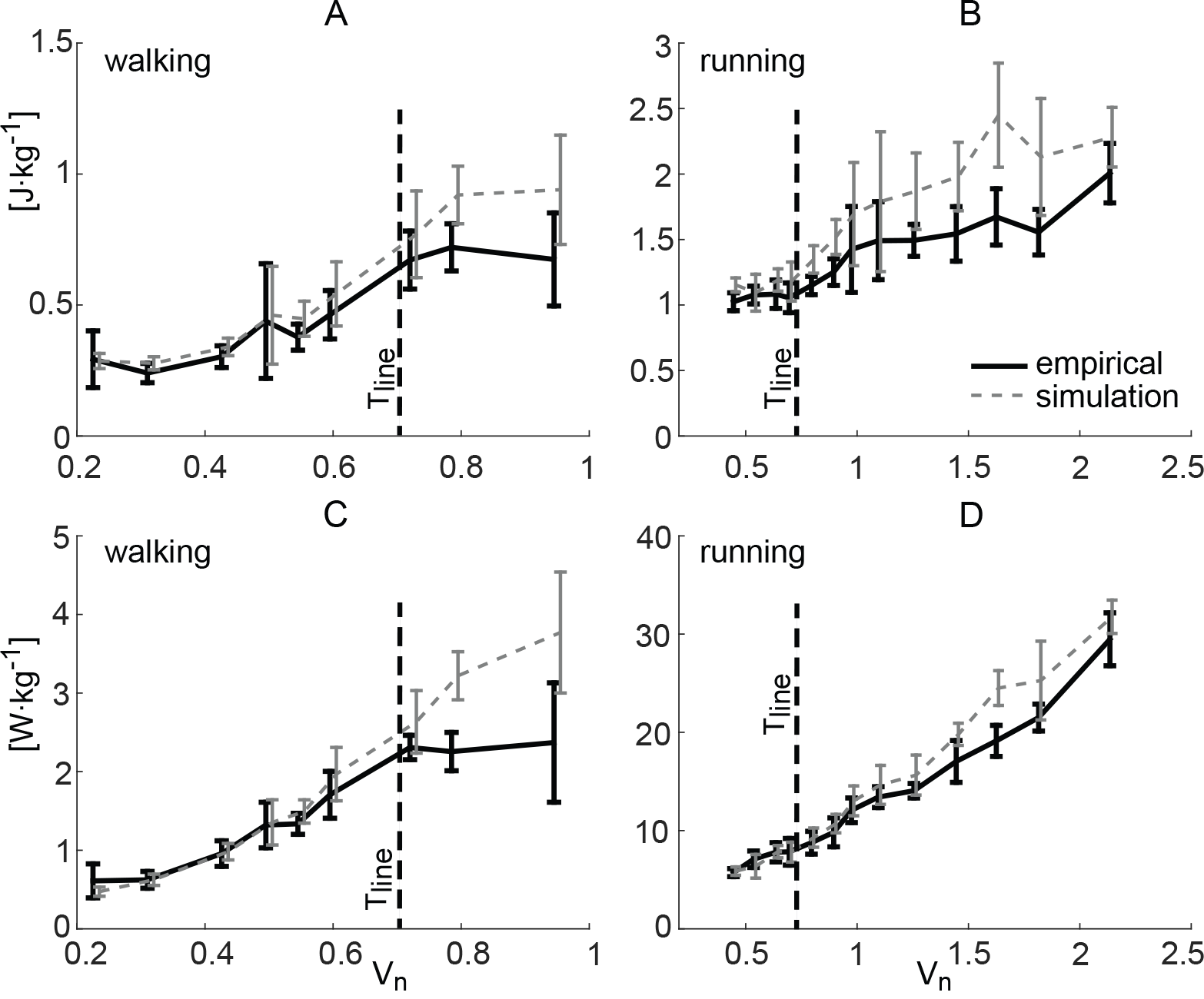
Empirical data (solid line) and simulation (dot-dashed line) of positive work per body mass (A – walking, B – running) and mean positive power per body mass (C – walking, D – running) during a step. Line represents mean value, error bar – one standard deviation. Vertical lines indicate conventional self-selected transition velocity between walking and running (T_line_).

### Protraction assumptions

While the weight-specific amplitude and shape of model limb vertical ground reaction force profiles are dependent on only duty factor (running) or duty factor, speed and leg length (walking), a further input is required to determine their duration (and impulse). This is achieved with assumptions concerning swing-leg (protraction) mechanics: we keep certain kinematic parameters constant in an effort to make protraction costs constant in order to assess only the implications of duty factor. Two assumptions are used; neither is expected to be completely valid, and their validity may change systematically with speed and/or gait.

Protraction assumption alternative 1: constant swing time T_sw_ (for a given speed). With this assumption, with increasing DF, step length and stance time increase too, but step frequency decreases. It suggests approximately constant effort for swinging the leg actively forwards (though not entirely as the angle swept during protraction does not remain constant). This assumption is possibly most appropriate in running due to the very active protraction (e.g. Weyand et al., 2000).

Protraction assumption alternative 2: constant step frequency F (for a given speed). With this assumption, with increasing DF, step length remains constant, stance time increases and swing time decreases. This assumption may be appropriate if aspects of both stance and swing leg mechanics are dominated by passive, gravity-driven motions (more likely in walking).

The values of the held-constant parameters are determined as a function of speed from regression equations taken from natural overground walking of our subjects:

- walking: 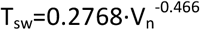, 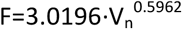,
- running: 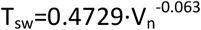, 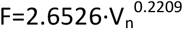.

### Generation of model parameter space

With the above model assumptions, the ground reaction forces, instantaneous power and the amount of positive mechanical work done can be calculated over a step. Numerical simulations were performed with normalized velocity 0.07-1.0 for walking and 0.25-2.6 for running (step 0.001) and with duty factor 0.505-0.75 for walking and 0.1-0.495 for running (step 0.0025); this bounds all experimental data. Simulation results are clearly not valid for all possible combinations of DF-V_n_. We assumed a maximum step length (for walking) and stance length (for running) of double leg length. Simulation results are also excluded (for very low speeds and large step lengths) for walking cases in which the centre of mass (COM) height is not at the mid-point of the stance; gaits with periods of negative horizontal velocity or limb tension were also excluded. The cost of transport (CT) space in terms of cost of muscle activation (C_MA_, as kilograms of active muscle mass [kg_am_]) is that which is more demanding between **m**ean **P**ower (*mP*_*cc*_) and positive **W**ork (*W*_*cc*_) during concentric contraction (per distance travelled – step length (SL) – and per body mass (BM), eq. 1):

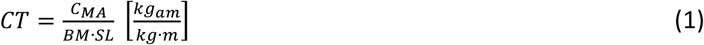

where

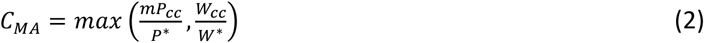

and P* and W* are muscle properties, described in next paragraph. Note that the power term is *not* cyclic or net-mean power; rather, it relates to the mean power demanded during the period of positive mechanical power – assumed to be during concentric muscle contraction (decoupling of these two periods by elastic mechanisms is assumed small and is neglected here).

### Initial assumed muscle properties

The cost of muscle activation following Hubel and Usherwood (2015) is based on muscle properties of 50 Joules work (*W\**) or 500 Watts power (*P\**) per kilogram of the active muscle mass (kg_am_). The ratio *W\**/*P\** equates to a muscle with V_max_=10 lengths s^−1^ operating at high power and efficiency (0.3 V_max_; Woledge et al., 1984) over reasonably high strain (30%). The ratio of these parameters gives a fundamental duration of 0.1s, and is based on empirical measurements of extreme work and cycle-power contractions. Deviation from these assumed muscle properties are considered further in the discussion.

### Empirical measurements

Walking and running were observed at various speeds in human subjects, initially allowing freely self-selected frequencies. The group consisted of 10 healthy adults (5 from Hubel and Usherwood, 2015, the same measurement technique and methodology) of a range of body dimensions (leg length range 0.87 – 1.01 m measured from ground wearing shoes to the greater trochanter; body mass range 55.1 – 95.3 kg). The study design was approved by the Royal Veterinary College ethics committee. All participants were volunteers and provided informed consent. At the beginning of the measurement session, the subject’s walking and running at preferred velocity was recorded; then the subject executed walk and run at speeds from ‘low’ to ‘high’ (an average of 5-6 trials for each type of locomotion).

A series of seven force platforms was used to record ground reaction forces (Kistler 9287B, sample frequency 500Hz), creating a 4.2m length and 0.9m width pathway, located in the central zone of the lab with free space for locomotion around 30m length. The data were averaged for each trail, of 2 to 7 consecutive steps. The duration of each phase, step frequency, duty factor, step length and velocity were calculated from forces and motions of centers of pressure. Power curves for each limb were calculated using the Individual Limb Method (Donelan et. al., 2002). From these, total positive work for each step, and the average power for the periods of positive power (again note – not mean cycle power), were calculated.

## Results

### Comparison of empirical and model mechanical work and power calculations

Model and empirical mean power for the period of positive work and total positive work match well for both walking and running (fig. 2) using appropriate measured kinematic data as model inputs. Disparity increases somewhat with speed. We take the match between model and measured data over the range of self-selected footfall timings (though enforced ‘walking’ or ‘running’ gaits) as support for the method for determining work (*W*_*cc*_) and mean power (*mP*_*cc*_) during muscle concentric contraction given unnatural footfall timings.

### Power, work and muscle activation cost surfaces

Model cost surfaces are shown for a range of cost functions, with the two alternative protraction timing assumptions (fig. 3). The three cost functions modelled are: positive work (J/(kg_bm_·m)) and mean power (W/(kg_bm_·m)) during concentric contraction; and the function we propose as relating closely to a metabolic cost function: the cost of muscle activation (kg_am_/(kg_bm_·m)), required from the more demanding between power and work (eq. 2).

**Figure 3.**
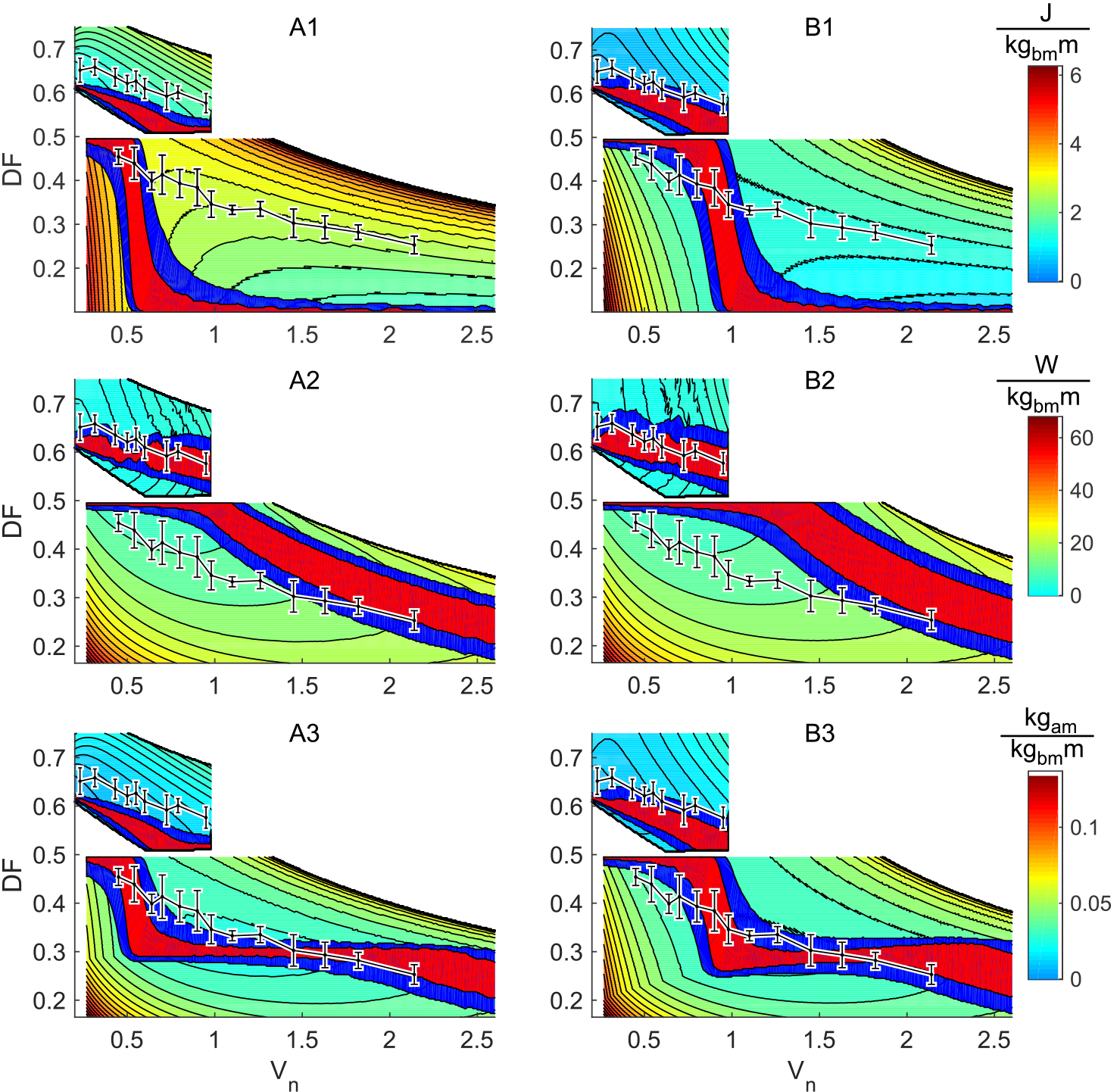
Cost of transport based on: positive work (1) and mean power (2) during concentric contraction, and muscle activation mass (3), for simulations with fixed swing time (A) and frequency (B). Red area denotes values with maximum range of 5% above minimum for given speed; blue shows 10% boundary. Data points show empirical mean (solid black line) and standard deviation (error bars). Results for walking are for DF above 0.5 and for running below this value.

Optimum DF for cost of transport based on positive work demand (CT_W_) at each velocity is similar for both gaits and both protraction models (fig. 3-A1, B1). The same is true for costs based on mean positive power demand (CT_P_) (fig. 3-A2, B2). However, simulations with constant frequency (fig. 3B) predict an optimum area somewhat closer to empirical DF. With initial assumed muscle properties (*W\**, *P\**), cost of transport, expressed as a muscle activation cost per distance per body mass (CT), is dominated by work demands during walking (CT_walk_=CT_Wwalk_; fig. 3-A3, B3), and a reasonable relationship (albeit with some offset) is predicted between speed and duty factor.

Optimal CT for running (fig. 3-A3, B3) is consistent with the work demand (CT_Wrun_) for low speeds, before a transition area where ‘cost of work’ and ‘cost of power’ surfaces cross (at points that match neither of their minimum values). From higher speed to the end of the range, power demand costs (CT_Prun_) were predicted to dominate. The predicted transition between work and power demand occurs at lower running speed with the constant swing time model (around 0.55 V_n_ vs 0.9 V_n_ for constant F model). At high velocities (above 1.4 V_n_) both models predict the same value of DF for minimum cost of transport.

### Comparison of model and inverse dynamic – derived ‘activation cost of transport’

Not only is the value of DF for a minimum of transport cost (fig. 3) close to empirical data, but also the predicted amount of costs obtained from the simulation matches those derived from inverse dynamics (fig. 4). For the assumed properties of muscle (*W\** = 50 J/kg_am_, *P\** = 500 W/kg_am_ and muscle concentric contraction duration = 0.1s), cost of transport during walking is much smaller than for running.

**Figure 4.**
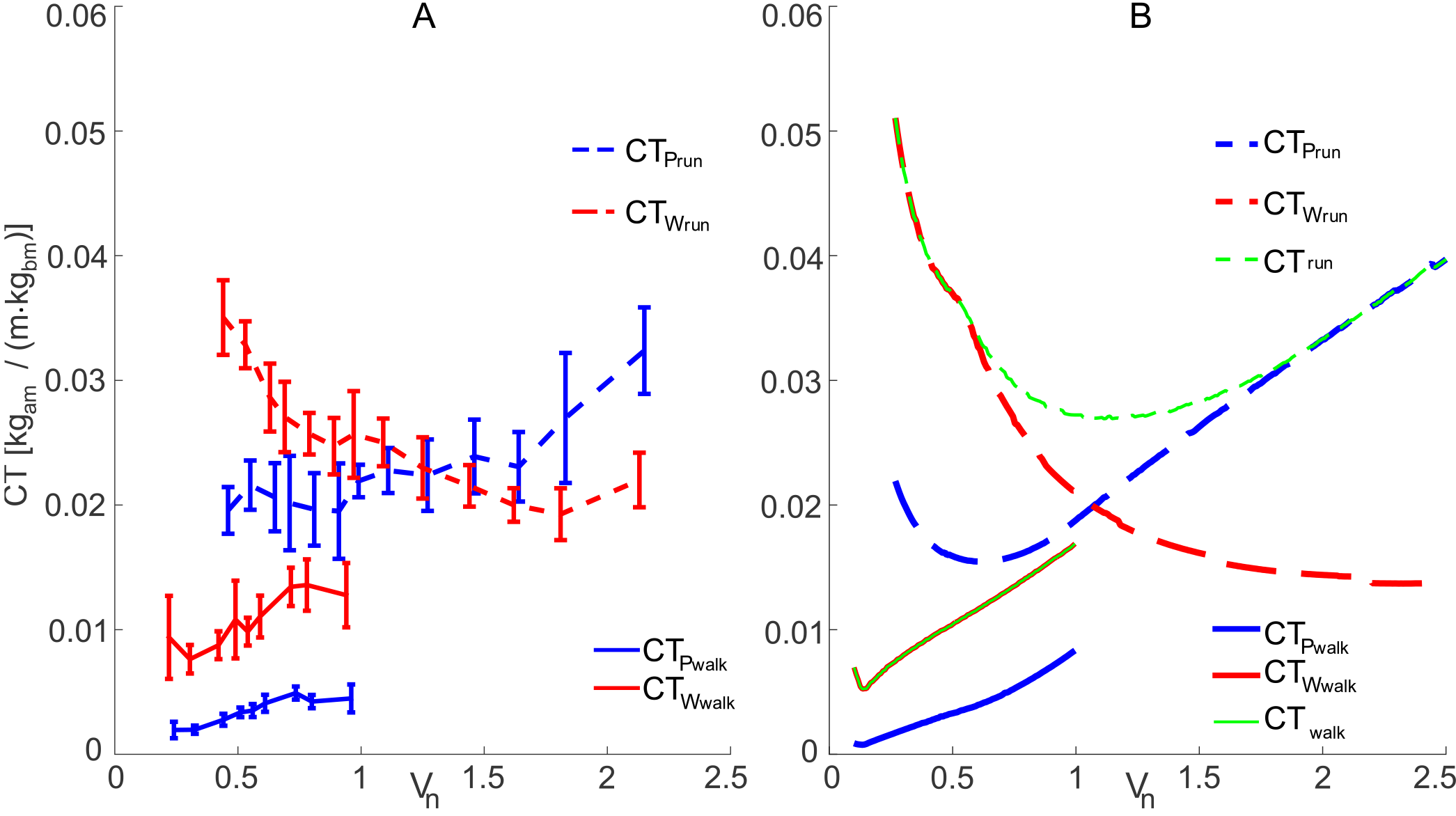
Empirical data (A) and predicted (B) cost of transport (green line) based on minimum muscle activation cost of walking (CT_walk_, solid line) and running (CT_run_, dash-dotted line). Blue lines denote power-demanded costs (CT_Pwalk_, CT_Prun_); red lines denote work-demanded costs (CT_Wwalk_, CT_Wrun_).

## Discussion

The presented results (specifically, fig. 3) provide support that the proposed muscle activation cost model relating to the demands of work and concentric contraction power provides a good account for the selection of duty factor with speed within both walking and running gaits. The current approach is based on a number of extreme assumptions. It assumes that the key muscle properties – *W\** and *P\** – are constant, and are in no way gait or speed dependent. Further, the initial values of these parameters – 50 J/kg and 500 W/kg, leading to a work: power ratio of 0.1s – are justified only as being ‘empirically reasonable’. Despite this, a good prediction of duty factor is made throughout the range of speeds of walking and running.

### Protraction models

The protraction models here are intended as two reasonable options for keeping the costs associated with swinging the leg forward each step approximately constant, enabling the focus of the modelling to be on the consequences of stance parameters on activation costs. The two protraction assumptions give similar simulation predictions, indicating that, while neither assumption necessarily succeeds in making swing costs exactly constant, the model predictions are not highly sensitive to this issue.

### Accounting for deviations from model predictions

The model predictions of duty factor are imperfect (slightly more or less so, depending on which protraction model is adopted), and the relative height of the walking and running surfaces are not satisfactory: walking is predicted to be less costly than running over the entire available speed range (Fig. 4c), including speeds well above the preferred walk-run transition and cross-over of reported metabolic cost of transports. This may be attributed – within the context of the proposed model – to two issues. The first is that the derivations of *mP*_*cc*_, the mean power during concentric contraction, may not be exactly comparable between walking and running due to their contrasting power profiles. The second is the assumption that the muscle properties are constant across gaits. Many details of the very different loading regimes of walking and running might well be expected to result in different muscle specific work and, especially, power capacities due, for instance, to recruitment of different ratios of muscle fibre types. Numerous other factors may account for deviation, from details of swing-leg mechanics to the numerous levels of additional complexity required for full consideration of both mechanics and muscle physiology. However, we proceed here by making small adjustments to the current model: namely, we tune the concentric contraction power capacity *P\** parameter in order to provide the closest fit to empirically measured duty factors within walking and running separately (maintaining *E\** at 50J/kg). Cost surfaces are shown (Fig. 5a,b) with *P\** tuned for each protraction model. Best-tuned conditions for walking are found with the constant frequency protraction assumption, and *P\** of 100W/kg; for running, the constant swing time protraction assumption is more successful, with a *P\** of 410W/kg. With these muscle parameters, the cost surfaces of walking raises with respect to running (compare Fig. 5c with Fig. 4b), resulting in a predicted walk-run transition speed based on activation cost minimization of 0.82 V_n_; better than the untuned model, but still somewhat high as the preferred transition speed is usually close to 0.70 V_n_. A further prediction of the low power capacity found in walking is the dominance of activation costs due to power over much of the speed range.

**Figure 5.**
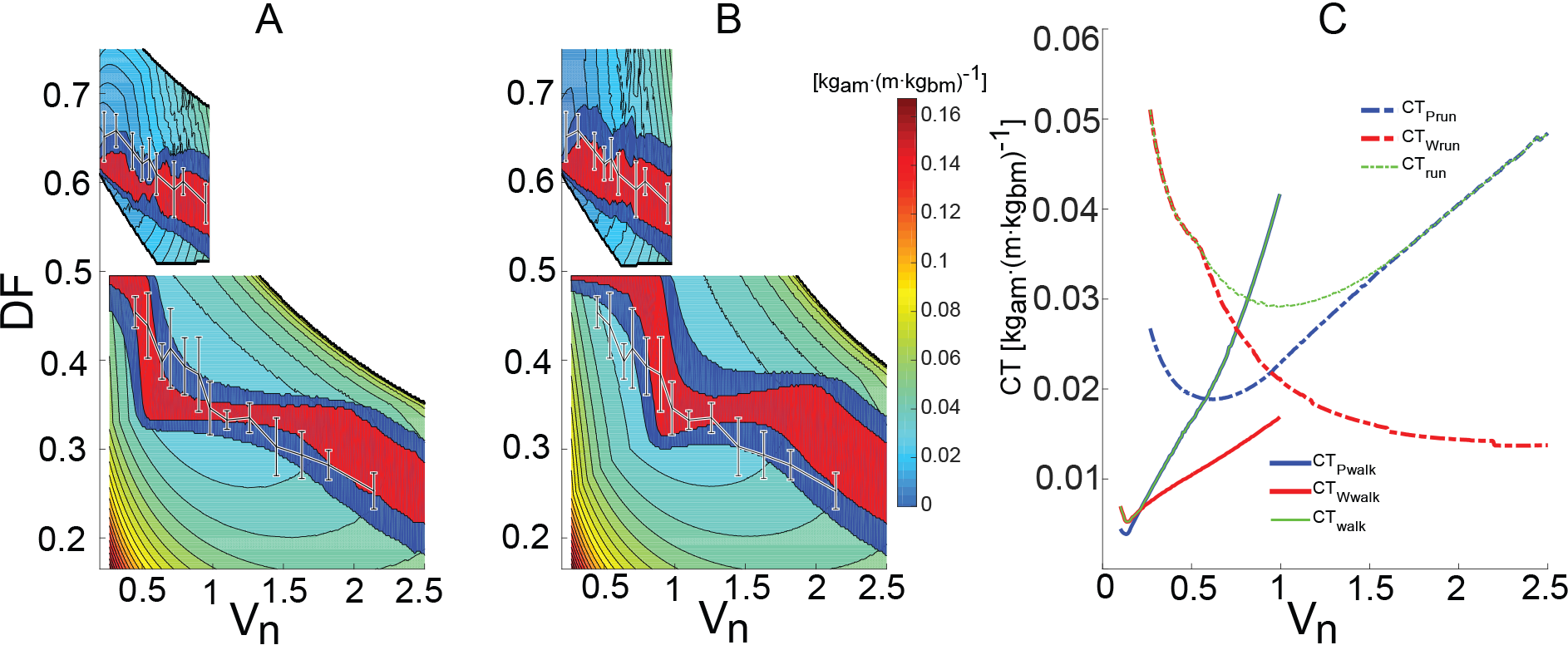
Cost of transport surfaces with tuned concentric contraction power capacities for both protraction models (A-constant T_sw_; B-constant F). Minimum values (C) across speed for best fit of real DF data to simulation results.

### Validating the tuned model

It is not surprising that a model with parameters tuned from the empirical data would have a better fit. For the model to have more than a descriptive value – to provide support that it represents an underlying mechanism – requires it to be used in predicting responses to some novel conditions. In order to achieve this, we model and measure responses to changes in step frequency at a range of speeds, and compare the predicted and observed duty factors.

The experimental conditions were with a range of velocities (moderate, lower and higher than moderate) and varied frequency (natural, lower and higher than natural) during overground walking. We used a transmission belt made of rope with markers stretched over rollers and driven by an electric motor to control locomotion velocity. Lower and higher frequencies were imposed by digital metronome and all subjects (N=5, body mass 73.8±4.4 kg, leg length 0.966±0.05 m) had the same conditions (velocities and non-natural frequencies). The results of model predictions (using tuned muscle parameters, and with both protraction assumptions) are shown together on one frequency – duty factor plane (fig. 6), by transforming the *T*_*sw*_ model results (eq. 3) onto that same plane with:

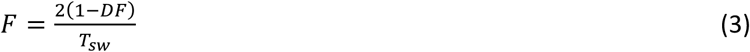

**Figure 6.**
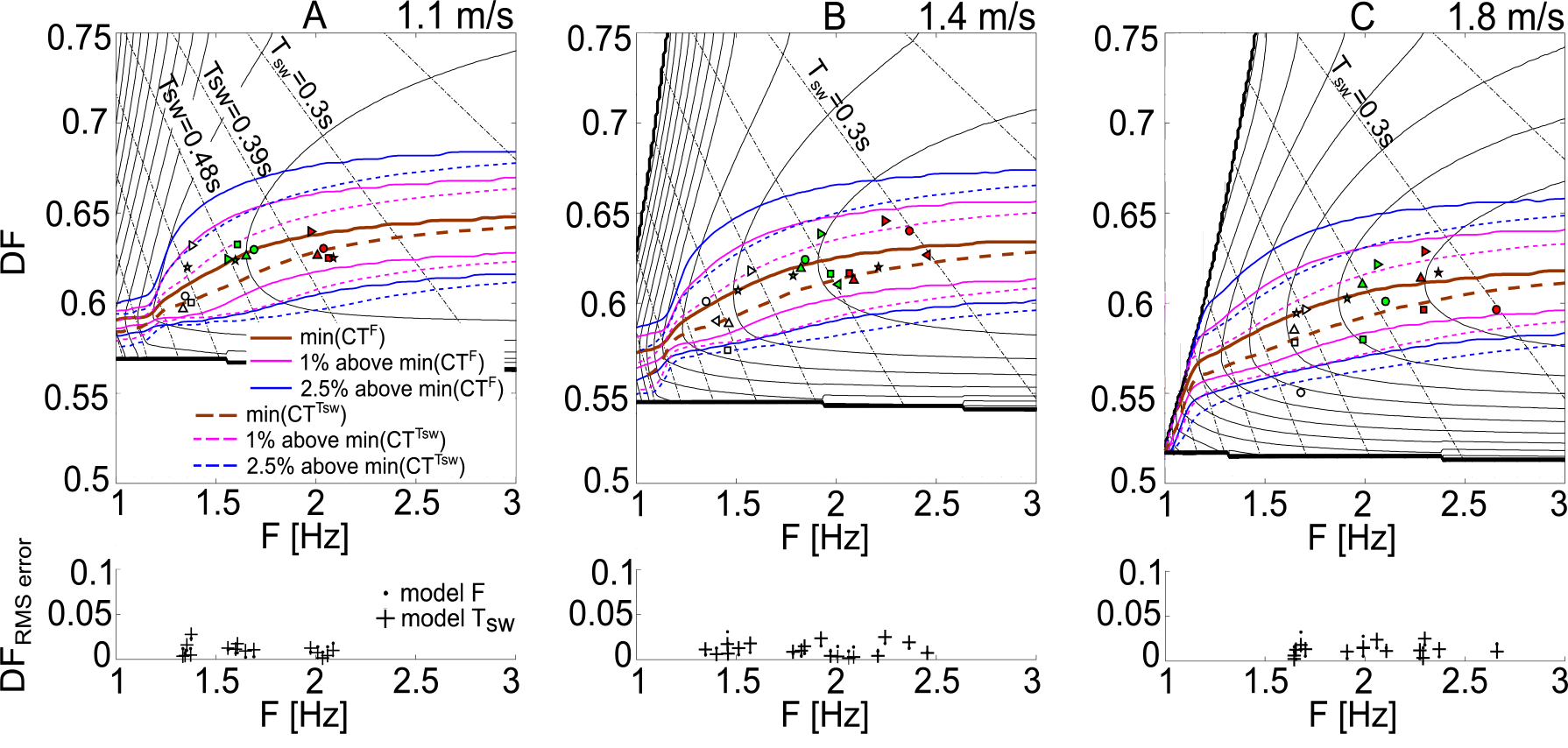
Cost of transport and prediction error (root mean square error) of constant frequency model (CT^F^, solid lines) and, transformed onto DF-F plane, the constant swing time model (CT^Tsw^, dashed lines). Black dash-dotted lines denote the iso-swing time’s contours. Observed results marked for walking with lower (white), higher (red) and preferred (green) step frequencies. Subjects denoted by marker shapes and results showed for low (A, 1.1 m/s), medium (B, 1.4 m/s) and high (C, 1.8 m/s) walking velocity. RMS error plots have results for both protraction models: dots for constant step frequency and crosses for constant swing time.

Models using the two protraction assumptions predict optimum solutions close to each other, with a little lower value of DF for the constant swing time assumption. Measurements of natural walking (green markers, fig. 6) unsurprisingly agree with the earlier observations and the simulation predictions using tuned muscle parameters. Responses to walking under non-natural conditions are successfully predicted from the modelled minimum cost of transport. White markers indicate measured duty factors for step frequencies lower, and red markers higher, than natural (figs. 6,7); these fall within 1% (for low velocity, fig. 6A) to 2.5% (for high velocity, fig. 6C) of the predicted minimum costs of transport. Prediction errors are comparable for the two models, but the constant frequency protraction assumption provided a marginally better prediction for non-preferred conditions.

**Figure 7.**
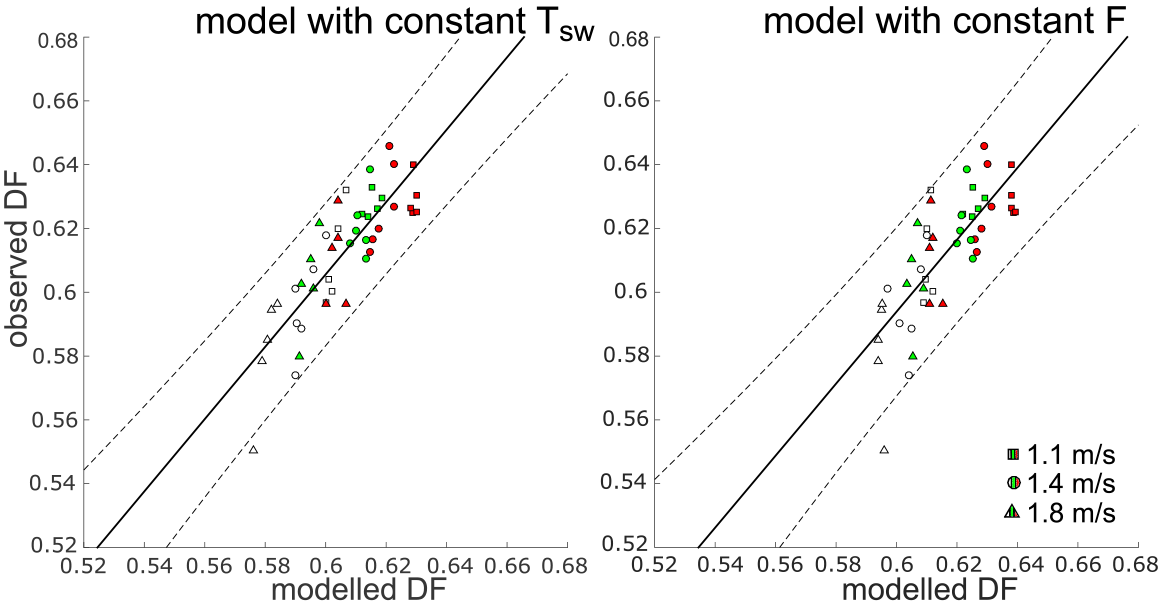
Observed vs modelled duty factor with confidence interval (95%) for three different walking velocities (squares – 1.1 m/s; circles – 1.4 m/s; triangles – 1.8 m/s) and three different walking conditions (step frequency, marker fill colour: white – low, green – natural and red – high).

## Conclusion

Minimization of a highly reductionist but mechanistic energetic cost model based on the costs of muscle activation due to mechanical work and power during contraction is broadly consistent with the duty factors observed in walking and running over a range of speeds. Further, the model has a successful predictive capacity in accounting for changes in walking duty factor under unnatural, high and low step frequency conditions. These findings provide further support that muscle activation during level locomotion may be the dominating cost, and that the ultimate demands for this activation may be related to work and power demands.

## Acknowledgments

We would like to thank our students, colleagues, friends and their relatives for providing walking and running data. We thank Zoe Self for her help in data collection and other people from Structure & Motion Lab for valuable comments. The work was funded by a Wellcome Trust Fellowship to JRU [095061/Z/10/Z]. RVC manuscript identifier: CBS_01321.

## Conflict of interest statement

The authors have no conflicts of interest.

## References

Alexander, R. McN. and Jayes, A. S., 1980. Fourier analysis of forces exerted in walking and running. J. Biomech. 13, 383–390.

Alexander, R. McN., 1980. Optimum walking techniques for quadrupeds and bipeds. J. Zool., Lond., 192, 97–111.

Alexander, R. McN., 1984. The gaits of bipedal and quadrupedal animals. Int. J. Robot. Res. 3, 49–59.

Alexander, R. McN., 2002. Energetics and optimization of human walking and running: the 2000 Raymond Pearl memorial lecture. American Journal of Human Biology, 14(5), 641–648.

Barclay, C. J., 2012. Quantifying Ca2+ release and inactivation of Ca2+ release in fast- and slow-twitch muscles. J. Physiol. 590, 6199–6212.

Bergmann, G., Graichen, F. and Rohlmann, A., 1993. Hip joint loading during walking and running, measured in two patients. J Biomech 26(8):969–90.

Bertram, J. E. A. and Ruina, A., 2001. Multiple walking speed-frequency relations are predicted by constrained optimization. J. Theor. Biol. 209(4), 445–453.

Biewener, A. A., 1983. Allometry of quadrupedal locomotion: The scaling of duty factor, bone curvature and limb orientation to body size. J. Exp. Biol. 105, 147–171.

Biewener, A. A., Farley, C. T., Roberts, T. J. and Temaner M., 2004. Muscle mechanical advantage of human walking and running: implications for energy cost. J. Appl. Physiol. 97(6): 2266–2274 doi: 10.1152/japplphysiol.00003.2004

Colbert, L. H., Hootman, J. M. and Macera, C. A., 2000. Physical activity-related injuries in walkers and runners in the aerobics center longitudinal study. Clin. J. Sport Med. 10(4):259–63.

Daley, M. A. and Usherwood, J. R., 2010. Two explanations for the compliant running paradox: reduced work of bouncing viscera and increased stability in uneven terrain. Biology Letters. 2010;6: 418–421. doi: 10.1098/rsbl.2010.0175.

Doke, J. and Kuo, A. D., 2007. Energetic cost of producing cyclic muscle force, rather than work, to swing the human leg. J. Exp. Biol. 210, 2390–2398.

Donelan, J. M., Kram, R. and Kuo, A. D., 2001. Mechanical and metabolic determinants of the preferred step width in human walking. Proc. R. Soc. Lond. B Biol. Sci. 268, 1985–1992.

Donelan, J. M., Kram R. and Kuo A. D., 2002. Simultaneous positive and negative external mechanical work in human walking. J. Biomech. 35: 117–124.

Farris, D. J. and Sawicki, G. S., 2012. The mechanics and energetics of human walking and running: a joint level perspective. J. R. Soc. Interface (2012) 9, 110–118 doi:10.1098/rsif.2011.0182

Gutmann, A. K., Jacobi, B., Butcher, M. T. and Bertram, J. E. A., 2006. Constrained optimization in human running. J. Exp. Biol. 209, 622–632 doi:10.1242/jeb.02010

Homsher, E., Mommaerts, W.F.H.M., Ricchiuti, N.V. and Wallner, A., 1972. Activation heat, activation metabolism and tension-related heat in frog semitendinosus muscles. J. Physiol. 220, 601–625.

Högberg, P., 1952. How do stride length and stride frequency influence the energy-output during running? Eur. J. Appl. Physiol. 14, 437–441 doi:10.1007/BF00934423

Hubel, T. Y. and Usherwood, J. R., 2015. Children and adults minimise activated muscle volume by selecting gait parameters that balance gross mechanical power and work demands. J. Exp. Biol. 218, 2830–2839 doi:10.1242/jeb.122135

Keller, T. S., Weisberger, A. M., Ray, J. L., Hasan, S. S., Shiavi, R. G. and Spengler, D. M., 1996. Relationship between vertical ground reaction force and speed during walking, slow jogging, and running. Clin. Biomech. 11(5):253–259.

Kram, R. and Taylor, C. R., 1990. Energetics of running: a new perspective. Nature 346, 265–267.

Lichtwark, G. A., Bougoulias, K. and Wilson, A. M., 2007. Muscle fascicle and series elastic element length changes along the length of the human gastrocnemius during walking and running. J. Biomech. 40 (2007): 157–164.

Margaria, R., Cerretelli, P., Aghemo, P. and Sassi, G., 1963. Energy cost of running. J. Appl. Physiol. 18, 367–370.

Mayhew, J. L., 1977. Oxygen cost and energy expenditure of running in trained runners. Br. J. Sports Med. 11, 116–121 doi:10.1136/bjsm.11.3.116

Nilsson, J. and Thorstensson, A., 1998. Ground reaction forces at different speeds of human walking and running. Acta Physiol. Scand. 136(2): 217–27.

Pontzer, H., Raichlen, D. A. and Sockol, M. D., 2009. The metabolic cost of walking in humans, chimpanzees, and early hominins. J. Hum. Evol. 56, 43–54.

Pontzer, H., 2016. A unified theory for the energy cost of legged locomotion. Biology Letters DOI: 10.1098/rsbl.2015.0935

Roberts, T. J., Kram, R., Weyand, P. G. and Taylor, C. R., 1998. Energetics of bipedal running. I. Metabolic cost of generating force. J. Exp. Biol. 201, 2745–2751.

Robilliard, J. J. and Wilson, A. M., 2005. Prediction of kinetics and kinematics of running animals using an analytical approximation to the planar spring-mass system. J. Exp. Biol. 208: 4377–4389; doi: 10.1242/jeb.01902

Schwartz, M. H., Rozumalski, A. and Trost, J. P., 2008. The effect of walking speed on the gait of typically developing children. J. Biomech. 41(8), 1639–1650 doi: 10.1016/j.jbiomech.2008.03.015

Selinger, J. C., O'Connor, S. M., Wong, J. D. and Donelan, J. M., 2015. Humans Can Continuously Optimize Energetic Cost during Walking. Curr. Biol. 2015 Sep 21;25(18):2452–6. doi: 10.1016/j.cub.2015.08.016. Epub 2015 Sep 10.

Sparrow, W. A. and Newell, K. M., 1998. Metabolic energy expenditure and the regulation of movement economy. Psych. Bull. Rev. 5, 173–196 doi:10.3758/BF03212943

Srinivasan, M. and Holmes, P., 2008. How well can spring-mass-like telescoping leg models fit multi-pedal sagittal-plane locomotion data? J. Theor. Biol. 2008 Nov 7; 255(1):1–7 doi: 10.1016/j.jtbi.2008.06.034. Epub 2008 Jul 15.

Srinivasan, M. and Ruina, A., 2006. Computer optimization of a minimal biped model discovers walking and running. Nature 439, 72–75.

Steudel-Numbers, K. L. and Wall-Scheffler, C. M., 2009. Optimal running speed and the evolution of hominin hunting strategies. J. Hum. Evol. 56, 355–360 doi:10.1016/j.jhevol.2008.11.002

Taylor, C. R., 1985. Force development during sustained locomotion: a determinant of gait, speed and metabolic power. J. Exp. Biol. 115, 253–262.

Usherwood, J. R. and Bertram, J. E. A., 2003. Gait transition cost in humans. Eur. J. Appl. Physiol. 90: 647–650 doi:10.1007/s00421-003-0980-6

Usherwood, J. R. and Hubel, T. Y., 2012. Energetically optimal running requires torques about the centre of mass. J. R. Soc. Interface. doi: 10.1098/rsif.2012.0145

Usherwood, J. R., Channon, A. J., Myatt, J. P., Rankin, J. W. and Hubel, T. Y., 2012. The human foot and heel-sole-toe walking strategy: a mechanism enabling an inverted pendular gait with low isometric muscle force? J. R. Soc. Interface 9, 2396–2402 doi:10.1098/rsif.2012.0179

Weyand, P. G., Sternlight, D. B., Bellizzi, M. J. and Wright, S., 2000. Faster top running speeds are achieved with greater ground forces not more rapid leg movements. J. Appl. Physiol. 89, 1991–1999.

Woledge, R. C., Curtin, N. A. and Homsher, E., 1984. Energetic aspects of muscle contraction. Monogr. Physiol. Soc. 41, 1–357.

Zarrugh, M. Y., Todd, F. N. and Ralston H. J., 1974. Optimisation of energy expenditure during level walking. European Journal of Applied Physiology and Occupational Physiology, 33 (4): 293–306

